# Mincle-GSDMD-mediated release of IL-1β containing small extracellular vesicles contributes to ethanol-induced liver injury

**DOI:** 10.1101/2022.11.30.518545

**Authors:** Quanri Zhang, Weiwei Liu, Katarzyna Bulek, Han Wang, Megan R. McMullen, Xiaoqin Wu, Nicole Welch, Renliang Zhang, Jaividhya Dasarathy, Srinivasan Dasarathy, Laura E. Nagy, Xiaoxia Li

## Abstract

**Background & Aims:** Macrophage inducible C-type lectin (Mincle) is expressed on Kupffer cells and senses ethanol-induced danger signals released from dying hepatocytes and promotes IL-1β production. However, it remains unclear what and how ethanol-induced Mincle ligands activate downstream signaling events to mediate IL-1β release and contribute to alcohol-associated liver disease (ALD). In this study, we investigated the association of circulating β-glucosylceramide (β-GluCer), an endogenous Mincle ligand, with severity of ALD and examined the mechanism by which β-GluCer engages Mincle on Kupffer cells to release IL-1β in the absence of cell death and exacerbates ALD.

**Approach and Results:** Concentrations of β-GluCer were increased in serum of patients with severe AH and correlated with disease severity. Challenge of Kupffer cells with LPS and β-GluCer induced formation of a *Mincle* and *Gsdmd*-dependent secretory complex containing chaperoned full-length GSDMD (Hsp90-CDC37-NEDD4) with polyubiquitinated pro-IL-1β and components of the Casp8-NLRP3 inflammasome loaded as cargo in small extracellular vesicles (sEV). Gao-binge ethanol exposure to wild-type, but not *Mincle*^*-/-*^ and *Gsdmd*^*-/-*^, mice increased release of IL-1β containing sEVs from liver explant cultures. Myeloid-specific deletion of *Gsdmd* similarly decreased the formation of sEVs by liver explant cultures and protected mice from ethanol-induced liver injury. sEVs collected from ethanol-fed wild-type, but not *Gsdmd*^*-/-*^, mice promoted injury of cultured hepatocytes and, when injected into wild-type mice, aggravated Gao-binge ethanol-induced liver injury.

**Conclusion:** β-GluCer functions as a DAMP activating Mincle-dependent GSDMD-mediated formation and release of IL-1β-containing sEVs, which in turn exacerbate hepatocyte cell death and contribute to the pathogenesis of ALD.

---

Alcohol-associated liver disease (ALD) ranges from steatosis to hepatitis, fibrosis, cirrhosis, and hepatocellular carcinoma(1). Severe alcohol-associated hepatitis (sAH) and chronic ALD are primary drivers of liver disease morbidity and mortality in the US, but effective treatment strategies are not available (2, 3). Inflammatory responses are critical contributors to progression of ALD. Impaired intestinal barrier integrity and changes in the microbiome contribute to increased circulating concentrations of microbes and their metabolites in patients with ALD and in animal models of ALD(4). Recognition of lipopolysaccharide (LPS) by Toll-like receptor 4 (TLR4) on resident hepatic macrophages (Kupffer cells) stimulates the expression of inflammatory cytokines, including TNFα and IL-1β. These inflammatory mediators in turn impact the functions of hepatocytes, liver sinusoidal endothelial cells and stellate cells, linking inflammation to loss of hepatocellular function and death (5-7). Multiple programmed cell death pathways are associated with ALD, including apoptosis, necroptosis and pyroptosis (8, 9). Caspase (Casp) 3, commonly associated with apoptosis, a relatively non-inflammatory form of cell death (10), is activated in livers of ethanol-fed mice and patients with AH (11, 12). Recent evidence suggests that Casp11-mediated pyroptosis also plays a fundamental role in ethanol-induced liver injury (13).

Abundant evidence indicates that the combination of increased circulating endotoxin and ethanol-induced hepatocellular death drives hepatic inflammation in ALD (1). However, the mechanisms by which the relatively low concentrations of endotoxin present in the context of alcohol consumption initiates chronic low-grade inflammation and how this is amplified in the progression of ALD are unknown. We reported that low concentrations of endotoxin, reflecting the relevant pathophysiological concentrations in both patients with ALD and ethanol-fed mice, induces expression of macrophage inducible C-type lectin (Mincle/Clec4e), a sensor for cell death, via IRAKM-dependent TLR4 signaling in hepatic macrophages (5). Mincle detects molecules released by dead hepatocytes, including β-glucosylceramide (β-GluCer), spliceosome-associated protein 130 (SAP130) and cholesterol sulfate, and activates inflammasomes and IL-1β production (14-16). Therefore, we proposed that Mincle serves as a critical link between cell death and inflammation in ALD (5).

IL-1β and IL-18 production by inflammasomes are critical drivers of hepatic inflammation and progression of ALD(1). The inflammasome is a Casp-containing multiprotein complex that processes pro-IL-1β and pro-IL-18 into their mature active forms (17-19). Mice deficient in inflammasome components, defective in IL-1β signaling, or provided with exogenous IL-1β receptor antagonist are protected from ethanol–induced liver injury (20, 21). IL-1β and IL18 concentrations are increased in patients with AH and are associated with disease severity (22-26). Two multicenter double blind, randomized, placebo controlled trials are currently evaluating the efficacy of Canakinumab (anti-IL-1β) (NCT037751090) or Anakinra (IL1 receptor antagonist)(NCT04072822) in AH.

Canonical IL-1β secretion involves initial processing of the inactive precursor of IL-1β by inflammasomes, followed by release of mature IL-1β from lytic cells (17). However, multiple non-canonical forms of non-lytic/cell death-independent release of IL-1β have also been described and are particularly important for export of mature IL-1β from neutrophils and macrophages in response to challenge with microbial products (27, 28). Human monocytes also release IL-1β in a Casp8-dependent alternative inflammasome activation pathway that is independent of pyroptosis (29). The contributions of non-canonical pathways of IL-1β release in the context of ALD are not well understood.

Gasdermin D (GSDMD) is classically associated with pyroptotic cell death, whereby GSDMD is cleaved by Casp1/11 and the N-terminal fragment of GSDMD then forms oligomeric pores in the plasma membrane, resulting in lytic cell death (30). Our recent study utilizing intestinal epithelial cells (IECs) revealed a novel form of IL-1β secretion mediated by a GSDMD-dependent non-pyroptotic release of small extracellular vesicles (sEVs). In IECs, we found that GSDMD is required for formation and release of sEVs containing polyubiquitinated pro-IL-1β upon activation of a Casp8-NLRP3 inflammasome (31). Mechanistically, full-length GSDMD is chaperoned by an Hsp90-CDC37 complex in IECs. In response to stimulus 1 (LPS), the chaperoned full-length GSDMD engages NEDD4, an E3 ubiquitin ligase, and brings it into the proximity of pro-IL-1β. The pro-IL-1β is then captured in the Casp8-NLRP3 inflammasome upon stimulus 2 (ATP). NEDD4 subsequently catalyzes the polyubiquitination of pro-IL-1β; polyubiquitination promotes the loading of the entire complex into vesicles destined for release into the extracellular space (31).

Since Mincle expressed on Kupffer cells senses ethanol-induced danger signals, contributing to IL-1β release and inflammatory responses (5), here we hypothesized that Mincle-dependent IL-1β production in Kupffer cells also relies on the GSDMD-dependent pathway we discovered in IECs. While in previous studies we identified SAP130 as an important DAMP activating Mincle in response to ethanol (5), here we report accumulation of another Mincle ligand, β-GluCer, in the circulation of patients with severe alcohol-associated hepatitis (sAH) and in mice after chronic ethanol feeding.

Both *Mincle* and *Gsdmd* were required for release of IL-1β-containing sEVs from hepatic macrophages and myeloid *Gsdmd-*deficient mice were protected from ethanol-induced liver injury. Provision of sEVs isolated from wild-type, but not *Gsdmd*-deficient, mice exacerbated ethanol-induced liver injury in mice. Taken together, these data identify β-GluCer-Mincle-GSDMD signaling in the regulation of IL-1β secretion in sEVs, providing an important link between hepatocellular injury and inflammation to drive the pathogenesis of ALD.

## Experimental Procedures

### Additional experimental details can be found in Supplemental Information Patient Samples

De-identified serum and plasma samples, along with basic clinical and demographic data, were obtained from the Northern Ohio Alcohol Center biorepository (NCT03224949). Patients with AH were stratified as moderate AH (MELD<20), or severe AH (MELD ≥20). Descriptive demographic and clinical data is provided in **Supplemental Table 1**. For western blots, samples from five livers explanted from patients with severe AH during liver transplantation and five wedge biopsies from healthy donor livers were snap frozen in liquid nitrogen and stored at -80°C. AH and healthy donor samples were provided by the NIAAA R24 Clinical Resource for Alcoholic Hepatitis Investigations at Johns Hopkins University. Clinical and demographic data on these subjects was previously reported (32). This study was approved by the Institutional Review Board at Cleveland Clinic (IRB 17-718) and all study participants consented prior to collection of data and blood samples.

### Mouse model

All mice were on C57BL/6 background. *Mincle* deficient, *Gsdmd* deficient and *Gsdmd*^*fl/fl*^ (Cyagen Biosciences) mice were previously described (5, 31, 33). Wild type mice were purchased from Jackson Laboratories. All procedures involving animals were approved by the Cleveland Clinic Institutional Animal Care and Use Committee. Ten- to twelve-week-old female knock out and heterozygous littermate mice were exposed to the Gao-binge (acute on chronic) model of ethanol exposure (34).

### Cell and liver explant culture

Primary hepatocytes and primary Kupffer cells from mice were isolated and cultured as previous described (5, 34). The immortalized mouse Kupffer cell line (imKC) was purchased from Sigma (Cat#SCC119). For liver explant culture, mouse livers were minced (∼2mm) and cultured overnight in serum-free culture medium (DMEM). Culture media were collected and used for isolation of EVs.

### Exosomes analysis

EVs were collected from culture medium of mice liver explant cultures or cultured cells. EVs were isolated using an exosome isolation kit (Invitrogen, 4478359) according to manufacturer’s instructions and subjected to nanoparticle tracking ZetaView analysis for quantification and sizing or for measurement of IL-1β by ELISA. The Polyethylene Glycol (PEG)-based density gradient method was used for enrichment of extracellular vesicles from human plasma.

### Data analysis and statistics

Data are expressed as mean ±SEM. For mouse feeding trials: n = 4 – 6 for Pair-fed, n= 6 - 8 for EtOH-fed. For cell culture experiments, at least 3 independent experiments were conducted. GraphPad Prism 7 was used for data analysis when Student’s t test was required, as well as for data representation. SAS (Carey, IN) was used for analysis of variance using the general linear models procedure and follow-up comparisons made by least square means testing. Data were log-transformed as necessary to obtain a normal distribution. p < 0.05 was considered significant.

## Results

### Serum β-glucosylceramide was increased in patients with AH and mice exposed to Gao-binge ethanol feeding

We recently reported that challenging peripheral blood mononuclear cells (PBMCs) from patients with sAH with low concentrations of LPS, equivalent to those detected in the circulation of patients with ALD, increases *Mincle* expression (35); secondary challenge with LPS and/or the Mincle ligand, trehalose-6,6-dibehenate (TDB), induced higher IL-1β expression in PBMCs from patients with sAH compared to healthy controls (35). β-Glucosylceramide (β-GluCer), an endogenous Mincle ligand released by damaged or dying cells, functions as an endogenous Danger Associated Molecular Pattern (DAMP) to amplify inflammatory responses (15), Circulating concentrations of β-GluCer are elevated in patients with diverse chronic inflammatory diseases, implicating β-GluCer as a functional biomarker for tissue damage and inflammation (36). Here we find that serum concentrations of the abundant d18:1/16:0 species of β-GluCer were increased in patients with severe AH and alcohol associated cirrhosis (AlcCir) compared with heathy controls (HC) (**Fig. 1A**). Model for end-stage liver disease (MELD) score, an indicator of severity of AH, was positively correlated with β-GluCer (d18:1/16:0) (**Fig. 1B**). The concentration of β-GluCer (d18:1/16:0) was also higher in plasma from mice after Gao-binge (Acute on chronic) ethanol feeding compared to pair-fed controls (**Fig. 1C**). Challenge of primary hepatocyte cultures with 150 mM ethanol for 24 h also increased the accumulation of β-GluCer in the culture media (**Fig. 1D)**.

**Figure 1.**
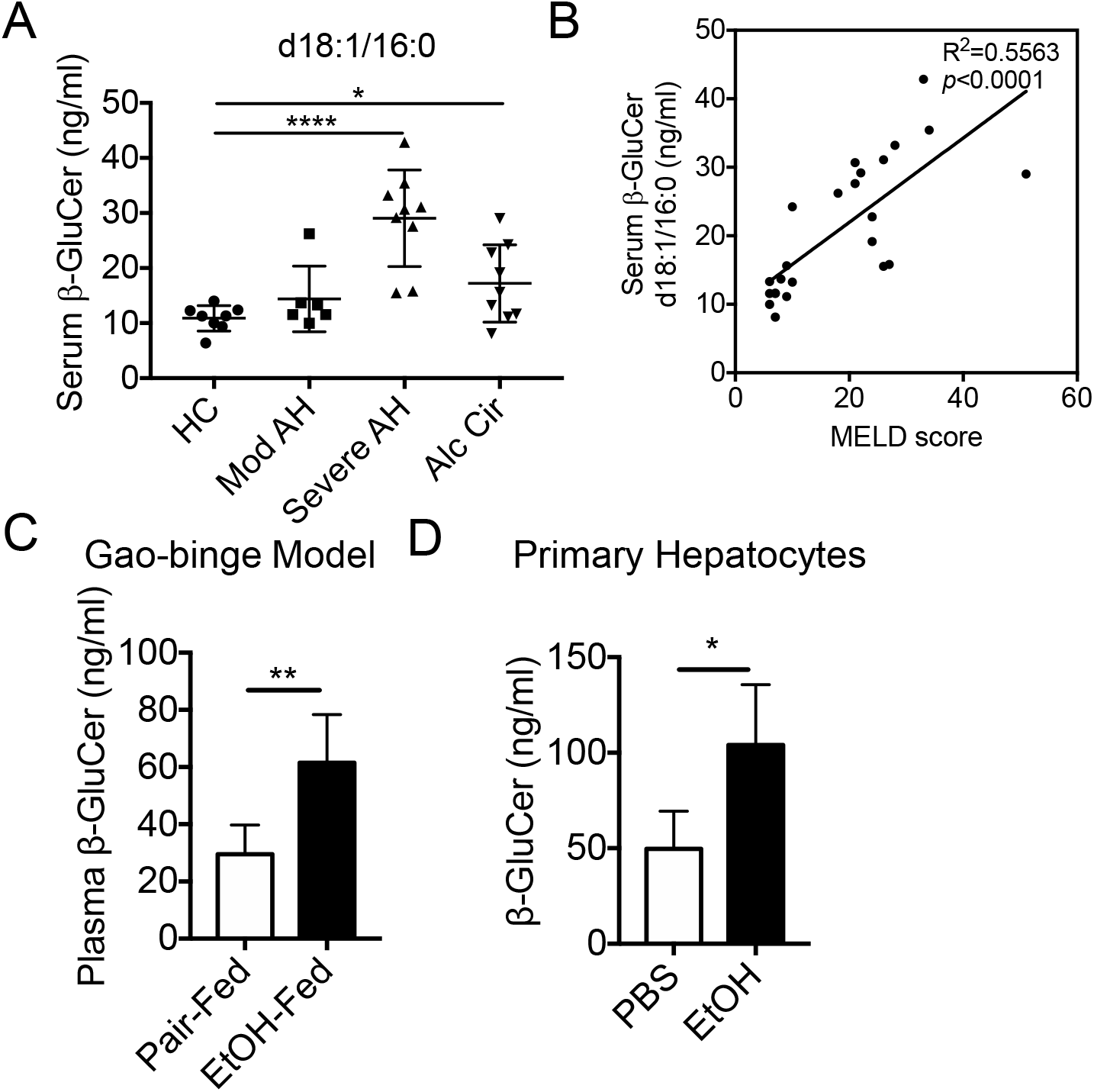
The Mincle ligand, β-GluCer, was increased in the circulation of patients with AH and mice after exposure to Gao-binge ethanol feeding. (**A**) Concentration of β-GluCer (d18:1/16:0) in serum of healthy controls (HC) (n=8) and patients with moderate AH (n=6), severe AH (n=9) or alcohol-associated cirrhosis (n=9). (**B**) Concentration of β-GluCer was correlated with MELD scores of patients with moderate and severe AH. (**C**) Concentration of β-GluCer in plasma of mice after Gao-binge ethanol feeding. (**D**) β-GluCer in culture medium of primary hepatocytes treated with 150mM EtOH for 24hrs. β-GluCer concentrations were measured by mass spectroscopy. Data represent mean ± SEM. ANOVA (A) or 2-tailed unpaired Student’s t test (C, D). *P<0.05, **P<0.01, ****P<0.0001.

### β-GluCer promoted Mincle-dependent IL-1β production without triggering cell death in Kupffer cells

We have previously reported that expression of Mincle on Kupffer cells is increased in response to chronic ethanol exposure and amplifies inflammatory responses in the liver by sensing the hepatocyte-derived danger signal SAP to promote IL-1β secretion (5). Since β-GluCer, another endogenous Mincle ligand with potent immunostimulatory activity (15), is elevated in patients with sAH and mice exposed to ethanol, we hypothesized that β-GluCer would also activate Mincle-expressing Kupffer cells. Primary cultures of Kupffer cells from *Mincle*^*+/-*^ and *Mincle*-deficient mice were first primed with a LPS (10ng/ml for 12h), and then stimulated with β-GluCer (20ug/ml for 12h) or ATP (1h), as a positive control. Pro-IL-1β expression in cell lysates was not affected by genotype or treatments (**Fig. 2A**). In contrast, β-GluCer stimulation increased IL-1β cleavage (**Fig. 2A**) and secretion into the cell culture media (**Fig. 2B**). The LPS/β-GluCer stimulated secretion of IL-1β was modest compared to that in LPS/ATP treated cells (**Fig. 2B**). Importantly, LPS/β-GluCer-stimulated, but not LPS/ATP-stimulated, IL-1β expression was dependent on *Mincle* (**Fig. 2A/B**). Intriguingly, whereas Kupffer cells treated with LPS/ATP produced IL-1β that was coupled with cell death, challenge of Kupffer cells with LPS/β-GluCer did not impact cell viability, assessed by LDH release (**Fig. 2C**). These data indicated that β-GluCer-activated Mincle signaling in Kupffer cells led to IL-1β secretion in the absence of lytic cell death.

**Figure 2.**
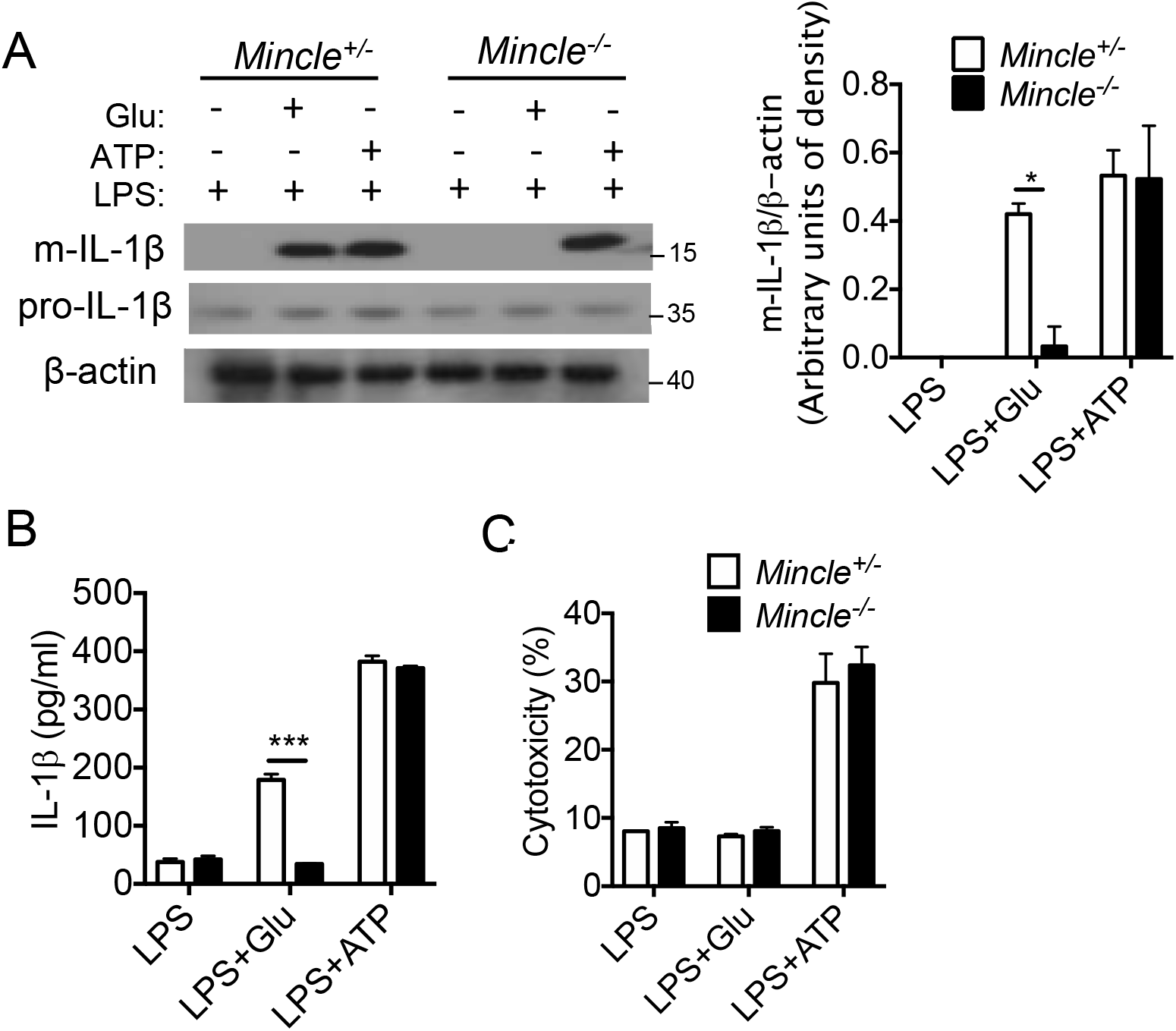
β-GluCer promoted Mincle-dependent IL-1β production without triggering cell death in Kupffer cells. Primary Kupffer cells isolated from *Mincle*^+/-^ and *Mincle*^-/-^ mice were primed overnight with 10 ng/ml LPS and then stimulated with 20ug/ml GluCer for 12 hrs or 2.5mM ATP for 1hr. **(A)** Cell lysates were collected for western blot analysis of pro- and mature-(m) IL-1β, with β-actin used as a loading control. **(B/C)** Cell culture medium was collected and used to (**B**) measure for IL-1β concentration by ELISA or (**C**) measure LDH release to assess cytotoxicity. Data represent mean ± SEM. 2-tailed unpaired Student’s t test. *P<0.05, ***P<0.001.

### Mincle-dependent IL-1β secretion was mediated by GSDMD-guided formation and release of small extracellular vesicles (sEVs)

We next investigated the mechanism for the non-lytic release of IL-1β in response to β-GluCer-activated Mincle signaling in Kupffer cells. Emerging evidence supports an important function for cell death-independent release of IL-1β (27-29), including our recent discovery of a non-pyroptotic role of GSDMD in the formation and release of IL-1β-containing sEVs from IECs (31). Therefore, we asked whether Mincle-dependent IL-β release from Kupffer cells is also achieved via activation of this GSDMD-mediated non-lytic pathway. To test this hypothesis, we knocked down *Gsdmd* (*Gsdmd*-KD) or *Mincle* (*Mincle*-KD) in an immortalized murine Kupffer cell line (imKC) using small hairpin RNA (shRNA). We primed wild-type, *Gsdmd* -KD and *Mincle*-KD imKCs with low concentration LPS (100pg/ml for 12h), followed by challenge with the Mincle ligands, β-GluCer (20ug/ml for 12h) or TDB (2ug/ml for 12h). Both LPS/β-GluCer and LPS/TDB induced a robust release of sEVs in wild-type imKCs; the size of the sEVs is illustrated by electron microscopy (**Fig. 3A**). Importantly, knock-down of *Gsdmd* or *Mincle* impaired the LPS/β-GluCer-and LPS/TDB-induced release of sEVs from imKCs (**Fig. 3A**).

**Figure 3.**
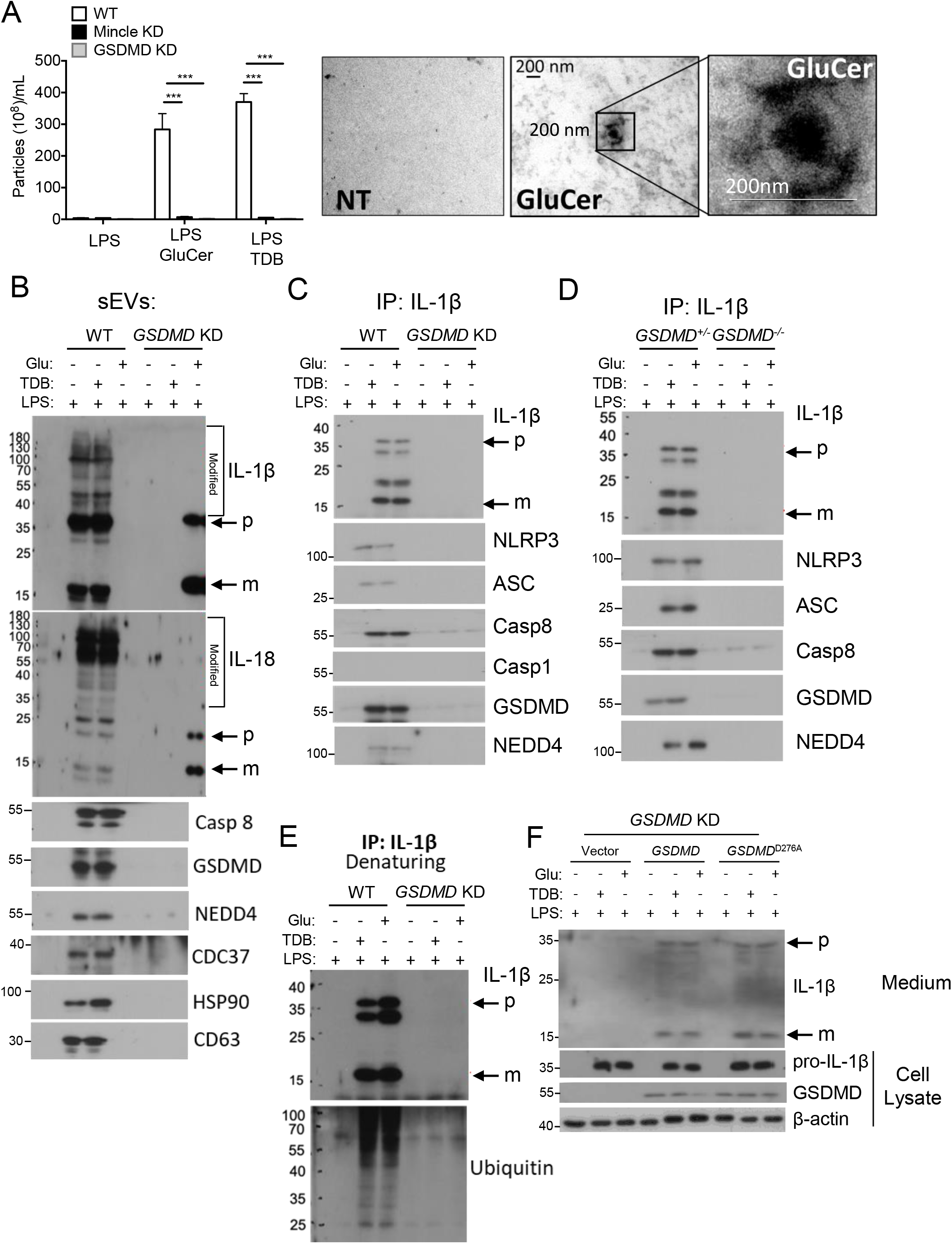
Mincle signal-dependent release of IL-1β containing sEVs from Kupffer cells requires GSDMD. **(A-D)** imKCs were transfected with shRNA to knockdown *Gsdmd* or *Mincle* and then primed overnight with 100 pg/ml LPS followed by stimulation with 20ug/ml GluCer or 2ug/ml TDB for 12 hrs. **(A)** sEVs released from immortalized murine KC (imKC) were quantified by ZetaView and visualized by electron microscopy. **(B)** Western analysis of sEVs from WT and *Gsdmd* KD imKCs utilizing antibodies against the indicated proteins (**Rec**: recombinant protein as + control; **p:** precursor; **m**: mature IL-1β or IL-18). *Note:* For all western using sEVs, sEVs isolated from an equal volume of cell culture media were loaded in gels; sEVs were not normalized for particle number. **(C)** IL-1β was immunoprecipitated from WT and *Gsdmd*-KD imKC culture media followed by western analyses with antibodies against the indicated proteins. **(D)** IL-1β was immunoprecipitated from the cell culture media from WT and *Gsdmd*-KD imKCs and probed with antibodies to IL-1β or Ubiquitin under denaturing conditions. **(E)** Primary KC isolated from heterozygous littermates and *Gsdmd*^*-/-*^ mice were primed overnight with 100 pg/ml LPS followed by stimulation with 20ug/ml GluCer or 2ug/ml TDB for 12 hrs. IL-1β was immunoprecipitated from culture media and analyzed by western blot as in **(C)**. (**F**)Expression vectors containing wild-type *Gsdmd* or D276A mutated *Gsdmd* were transfected into *Gsdmd*-KD imKCs, then primed with 100 pg/ml LPS overnight and stimulated with 20ug/ml GluCer or 2ug/ml TDB for 12 hrs. Cell lysates and culture medium were collected for western analyses for indicated proteins. N=3 independent experiments. Data represent mean ± SEM. One way ANOVA **P<0.01, ***P<0.001.

We have previously reported that when the Casp8 inflammasome is activated in intestinal epithelial cells (IECs), full-length GSDMD is chaperoned by CDC37/HSP90 and recruits NEDD4 (an E3 ligase) to the complex (31). NEDD4 in turn ubiquitinates pro-IL-1β; this facilitates the loading of pro-IL-1β into the cargo of the sEVs (31). Importantly, sEVs released from LPS/β-GluCer-treated imKCs contained higher molecular weight modified pro-IL-1β, pro-Il-18, Casp8, full-length GSDMD, NEDD4, HSP90 and CDC37 (**Fig. 3B**). The IL-1β/IL-18 containing sEVs from LPS/β-GluCer treated imKCs were also positive for the exosome marker CD63 (**Fig. 3B**). Taken together, these data are consistent with the hypothesis that β-GluCer-activated Mincle signaling in Kupffer cells utilizes the GSDMD-mediated non-lytic pathway to release IL-1β/IL-18 containing sEVs.

While the sEVs contained both IL-1β and IL-18 inflammasome products, we focused our mechanistic studies on IL-1β. When IL-1β was immunoprecipitated from the culture media of LPS/β-GluCer-treated imKCs (**Fig. 3C)** or primary Kupffer cells (**Fig. 3D**), these same GSDMD-interacting partners were present in a secretory complex along with pro- and mature-IL-1β and Casp8-NLRP3 inflammasome components (**Fig. 3C/D**).

Importantly, *Gsdmd* was required for both the release of sEVs containing IL-1β, IL-18, GSDMD, NEDD4, and Casp8 (**Fig. 3B**) and the release of the IL-1β secretory complex (**Fig. 3C/D**). Lysates from the *Gsdmd-*KD imKCs and primary Kupffer cells from *Gsdmd-/-* mice had similar expression of pro-IL-1β, Casp8 and NEDD4 (**Sup. Fig. 2A/B**). Finally, given the presence of the E3 ligase NEDD4 in both the sEVs and secretory complex from wild-type Kupffer cells, we asked whether the higher molecular weight/modified forms of IL-1β in the sEVs were poly-ubiquitinated. Immunoprecipitates of IL-1β from the media of wild-type, but not *Gsdmd-*KD, imKCs, performed under denaturing conditions, had more ubiquitin immunoreactivity in response to LPS/β-GluCer and LPS/TDB (**Fig. 3E)**. Taken together, our results indicate that signaling by Mincle ligands mediates IL-1β secretion via the release of GSDMD-dependent CD63^+^ sEVs from Kupffer cells.

Pyroptotic activation of GSDMD requires cleavage of D276, in the linker region of GSDMD, in response to inflammasome assembly (37). Since β-GluCer-activated Mincle signaling in Kupffer cells led to IL-1β secretion in the absence of lytic cell death (**Fig. 2C**), we hypothesized that expression of the inactive D276A form of GSDMD in *Gsdmd*-KD imKCs would restore their ability to release IL-1β-containing sEVs. Indeed, when *Gsdmd*-KD imKCs were transduced with either wild-type or GSDMD^D276A^, both LPS/β-GluCer and LPS/TDB treatment stimulated the release of IL-1β into the culture media (**Fig. 3F**).

### *Mincle*- or *Gsdmd*-deficiency impaired the release of IL-1β-containing sEVs from liver explants from mice after Gao-binge ethanol feeding

Since both *Mincle-/-* and *Gsdmd-/-* were required for the release of IL-1β-containing sEVs from Kupffer cells, we next explored their role in the release of sEVs from mouse liver. *Mincle-/-, Gsdmd*-/-, and their respective heterozygous littermates were subjected to Gao-binge ethanol feeding. Liver explants were cultured overnight and exosomes were isolated from the culture media by standard methods. The size distribution of sEVs released by liver explants was not affected by genotype or ethanol feeding (**Fig. 4A/B**). In contrast, Gao-binge ethanol increased the number of sEVs and their IL-1β cargo released from liver explants from heterozygous littermates, but not *Mincle-/-* or *Gsdmd*-/-, mice (**Fig. 4C/D**). In order to characterize the composition of the secretory complexes released from *Gsdmd*^*+/-*^ and *Gsdmd*^*-/-*^ mice, we normalized total sEV numbers released from liver explants between genotypes and immunoprecipitated IL-1β. Gao-binge ethanol feeding induced the formation of a secretory complex containing full-length GSDMD (Hsp90-CDC37-NEDD4) with high molecular weight modified pro-IL-1β and Casp8 in wild-type, but not *Gsdmd-/-*, mice (**Fig. 4E**). We next asked if extracellular vesicles circulating in patients with moderate and sAH also contained modified high molecular weight IL-18. Indeed, exosomes isolated from patients with AH contained pre-dominantly higher molecular weight forms of IL-18 (**Fig. 4F**). Taken together, these data indicate that Gao-binge ethanol triggers a Mincle-GSDMD-dependent release of CD63+ IL-1β-containing sEVs.

**Figure 4.**
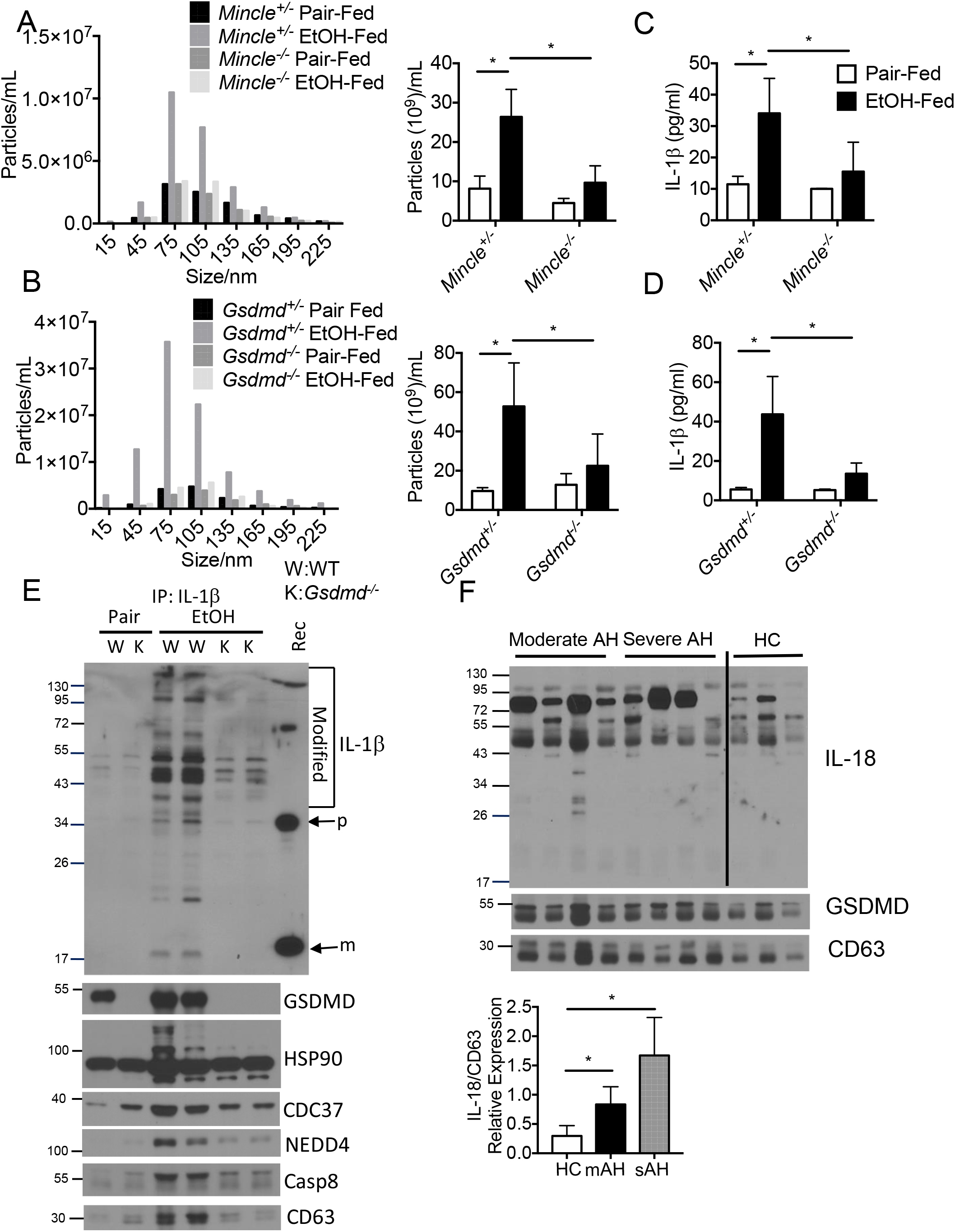
Mincle and GSDMD promoted IL-1β secretion via small extracellular vesicles (sEVs) **(A-E)** *Gsdmd*^-/-^ or *Mincle*^*-/-*^ and heterozygous littermate mice (n=6) were exposed to Gao-binge ethanol feeding. **(A**,**B)** 100 mg of liver explants were cultured overnight. sEVs were isolated from the cell culture media and analyzed by nanoparticle tracking with ZetaView for size distribution and total EV numbers/ml media. **(C**,**D)** Concentration of IL-1β in the collected sEVs was measured by ELISA. (**E**) IL-1β was immunoprecipitated from culture medium for western blotting with the indicated antibodies. Recombinant IL1-β (**Rec**) was included as a positive control. **p**: precursor; **m**: mature IL-1β; **W**: wild type; **K**: knock-out). **(F)** EVs isolated from plasma of healthy controls (HC) and patients with AH were analyzed by western blotting with antibodies against the indicated proteins. Data represent mean ± SEM. ANOVA *P<0.05, **P<0.01, ***P<0.001.

### Myeloid *Gsdmd*-deficiency protects mice from Gao-binge induced liver injury

Consistent with the hypothesis that the Mincle-GSDMD-mediated release of IL-1β-containing sEVs in response to ethanol contributes to the progression of ethanol-induced liver injury, we have previously reported that *Mincle-/-* are protected from chronic ethanol-induced liver injury (5). Similarly, GSDMD has been implicated in progression of ethanol-induced liver injury (13, 38). Hepatocyte overexpression of a constitutively active GSDMD exacerbated liver injury (13) and global *Gsdmd-/-* mice are protected from Gao-binge induced liver injury (38). We also confirmed by global *Gsdmd-/-* mice are protected from Gao-binge ethanol with reduced circulating ALT, hepatic triglycerides and inflammatory cytokine expression (**Suppl. Fig. 2**). While these previous studies have focused on GSDMD activity in hepatocytes, our data on GSDMD activity in Kupffer cells led us to hypothesize the myeloid *Gsdmd*-deficiency would be critical for the formation of IL-1β sEVs and the development of ethanol-induced liver injury. We generated LysM-Cre *Gsdmd*^*fl/fl*^ mice to test this hypothesis. As expected, *Gsdmd*^*fl/fl*^ mice developed the typical profile of Gao-binge induced liver injury, including increased circulating ALT, hepatic steatosis and increased expression of inflammatory cytokines and serum amyloid A, an acute phase protein (**Fig. 5A-D**). However, myeloid-*Gsdmd*-deficient mice were protected from Gao-binge induced liver injury (**Fig. 5A-D**). Importantly, while the number of sEVs released from liver explants was not affected by genotype (**Fig. 5E**), the IL-1β content in sEVs was reduced in myeloid-*Gsdmd*-deficient mice (**Fig. 5F**)

**Figure 5.**
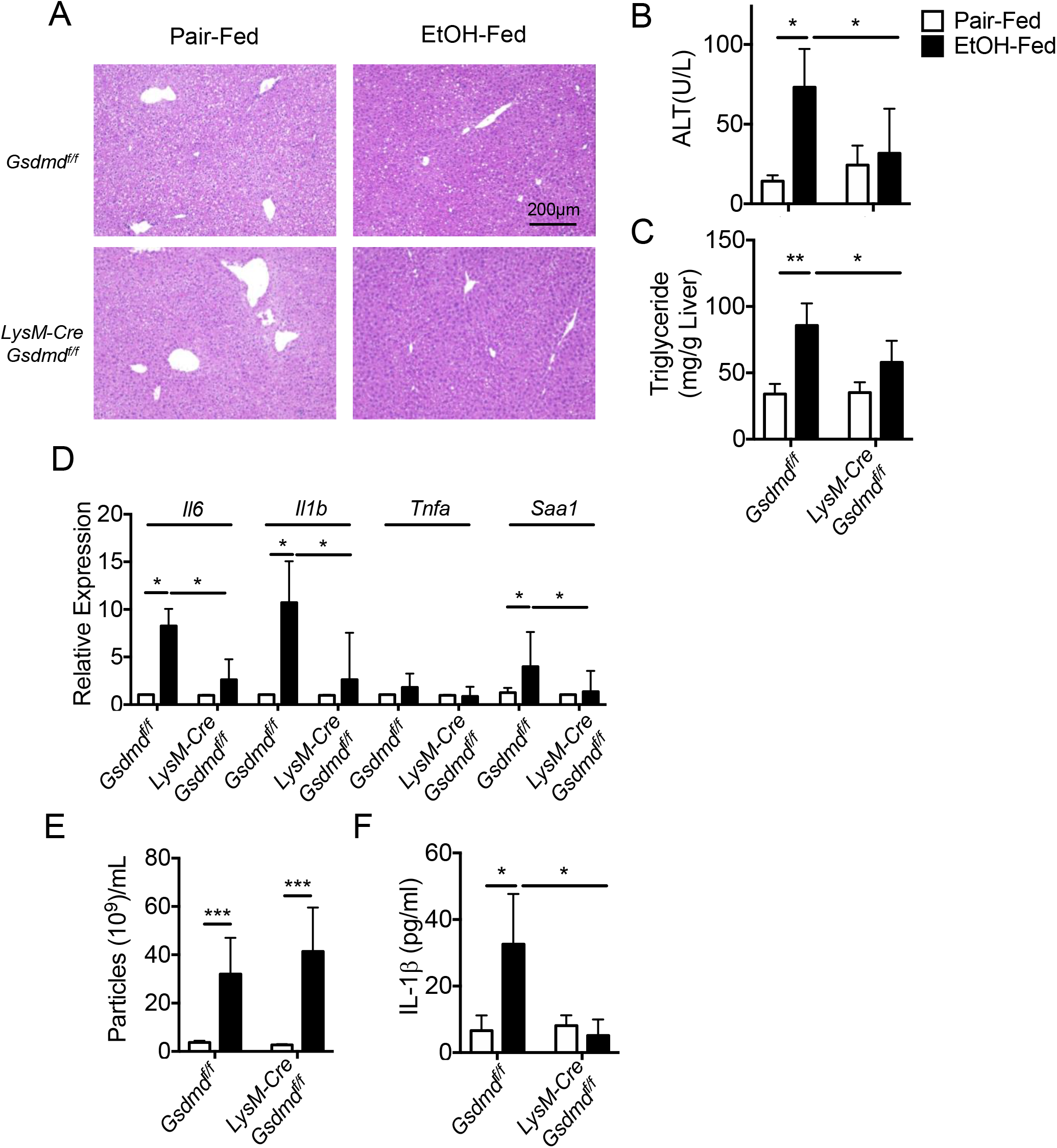
Myeloid GSDMD promoted release of IL-1β containing sEVs. *LysM-Cre Gsdmd*^f/f^ and *Gsdmd*^f/f^ mice (n=6) were exposed to Gao-binge ethanol feeding. (**A**) H&E staining for liver histology. (**B**) ALT activity in plasma. **(C)** Hepatic triglyceride content. (**D**) IL-6, IL-1β, TNFα and SAA1 mRNA expression. **(E)** sEVs from liver explants cultures were quantified by ZetaView as described in Fig 4. **(F)** sEV numbers were equalized and the concentration of IL-1β measured by ELISA. Data represent mean ± SEM. ANOVA. *P<0.05, **P<0.01, ***P<0.001.

### IL-1β-containing sEVs promote hepatocyte death and liver injury

Hepatocytes are highly responsive to cytokines/chemokines produced by Kupffer cells, such as IL-1β, contributing to the pathogenesis of ALD. We recently reported that IL-1β-induced expression of the acute phase protein SAA1; this response, in combination with ethanol, increased Casp3 cleavage and hepatocyte death (34). These data led us to hypothesize that IL-1β-containing sEVs may exert their pathogenic role, at least in part, via induction of the acute phase protein SAA1. As expected, challenge of hepatocytes with ethanol or recombinant IL-1β induced expression of SAA and cytotoxicity (**Fig. 6A/B**)(34). Importantly, IL-1β-containing sEVs isolated from liver explant cultures of wild-type mice after Gao-binge ethanol exposure also induced the expression of SAA1 and death of primary hepatocytes (**Fig. 6A/B**).

**Figure 6.**
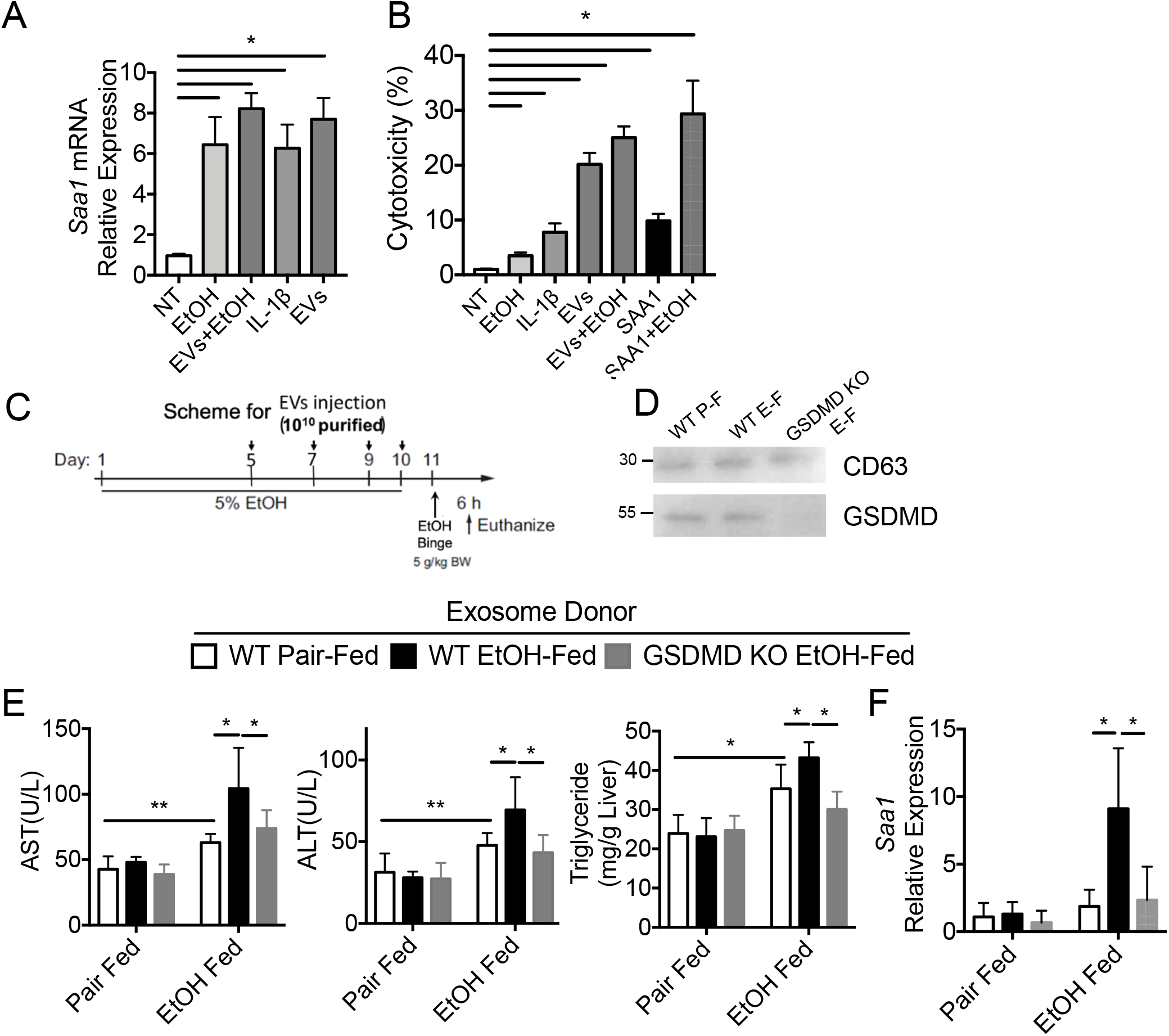
Exosomes from liver explant cultures of Gao-binge mice promote liver injury. (**A, B**) Primary hepatocytes were treated with recombinant IL-1β (10 ng/ml) or sEVs isolated from liver explant cultures of Gao-binge EtOH-fed WT mice. **(A)** Expression of SAA1 mRNA. **(B)** Cytotoxicity was assessed by LDH assay. **(C-F) (C)** Schematic diagram illustrating the protocol for exosome donor experiment. sEVs were isolated from liver explant cultures from WT Pair-fed, WT EtOH-fed and *Gsdmd-/-* EtOH-fed mice. WT mice were injected with 10^10^ sEVs during Gao-binge acute on chronic ethanol feeding. (**D**) sEV numbers were normalized and CD63 and GSDMD assessed by western blot. (**E**) AST/ALT activity in plasma and hepatic triglyceride content. **(F)** SAA1 mRNA expression. Data represent mean ± SEM. ANOVA. *P<0.05, **P<0.01, ***P<0.001.

These cell culture data suggest that IL-1β-containing sEVs might contribute to chronic ethanol-induced liver injury. Therefore, to determine the *in vivo* biological activity of sEVs, sEVs were collected from liver explants from wild-type and *Gsdmd-/-*mice and 10^10^particles were injected intraperitoneally to wild-type mice during the course of Gao-binge ethanol feeding (**Fig. 6C**). Expression of CD63 was equal across the purified sEVs, but GSDMD was absents from the sEVs isolated from *Gsdmd-*deficient mice (**Fig. 6D**). Importantly, sEVs purified from liver explant cultures of wild-type ethanol-fed, but not pair-fed, mice markedly increased hepatic injury, steatosis and expression of inflammatory cytokines and SAA (**Fig. 6E-F**). This exacerbation was not observed with sEVs purified from liver explant cultures of *Gsdmd-/-* ethanol-fed mice (**Fig. 6E-F**). Taken together, these results suggest that GSDMD-mediated production of IL-1β-containing sEVs contribute to the pathogenesis of ALD.

## Discussion

In this study, we report Mincle-dependent IL-1β secretion via GSDMD-guided formation and release of sEVs from Kupffer cells during ethanol-induced liver injury. This pathway is triggered by β-glucosylceramide (β-GluCer), an endogenous Mincle ligand released by dying hepatocytes after ethanol exposure. Importantly, serum β-GluCer is elevated in patients with AH and positively correlates with disease severity. β-GluCer is also elevated in the circulation of mice exposed to Gao-binge ethanol feeding. *Gsdmd*- or *Mincle*-deficiency impaired the release of IL-1β containing sEVs and IL-1β containing sEVs exacerbated hepatocyte cell death. Intravenous injection of IL-1β containing sEVs purified from liver explant cultures of ethanol-fed, but not pair-fed, mice markedly increased ethanol-induced hepatic injury and steatosis, indicating that IL-1β containing sEVs contribute to the pathogenesis of ALD.

Previous studies have suggested that Mincle ligands induce activation of the ASC-NLRP3 inflammasome, which leads to Casp8-dependent IL-1β production (5, 39). In this study, we showed that the Mincle ligand β-GluCer induces GSDMD and its interacting partners (including E3 ligase NEDD4) to form a secretory complex with the Casp8-inflammasome and polyubiquitinated pro-IL-β (**Fig. 7**). This secretory complex is loaded into sEVs, which are marked by the exosome marker CD63. We recently reported the downstream signaling events of Casp8-inflammasome activation in IECs, where full-length GSDMD, chaperoned by CDC37/HSP90, recruits NEDD4 to mediate polyubiquitination of pro-IL-1β, followed by cargo loading into CD63^+^ sEVs (31). Pyroptotic activaton of GSDMD is triggered via cleavage at D276, located in its linker region, by either Casp1 or Casp8 in response to inflammasome assembly. β-GluCer-activated Mincle signaling in Kupffer cells led to the release of IL-1β-containing sEVs in the absence of cytotoxicity. Importantly, when *Gsdmd*-deficient imKCs (**Fig. 4F**) or IECs (31) are transduced with either wild-type or GSDMD^D276A^ mutant, release of IL-1β-containing sEVs is restored, demonstrating a non-lytic function of GSDMD in the process of sEV release. Taken together, these results indicate that β-GluCer-activated Mincle signaling in Kupffer cells utilizes a non-lytic GSDMD-mediated mechanism to release IL-1β-containing sEVs.

**Figure 7.**
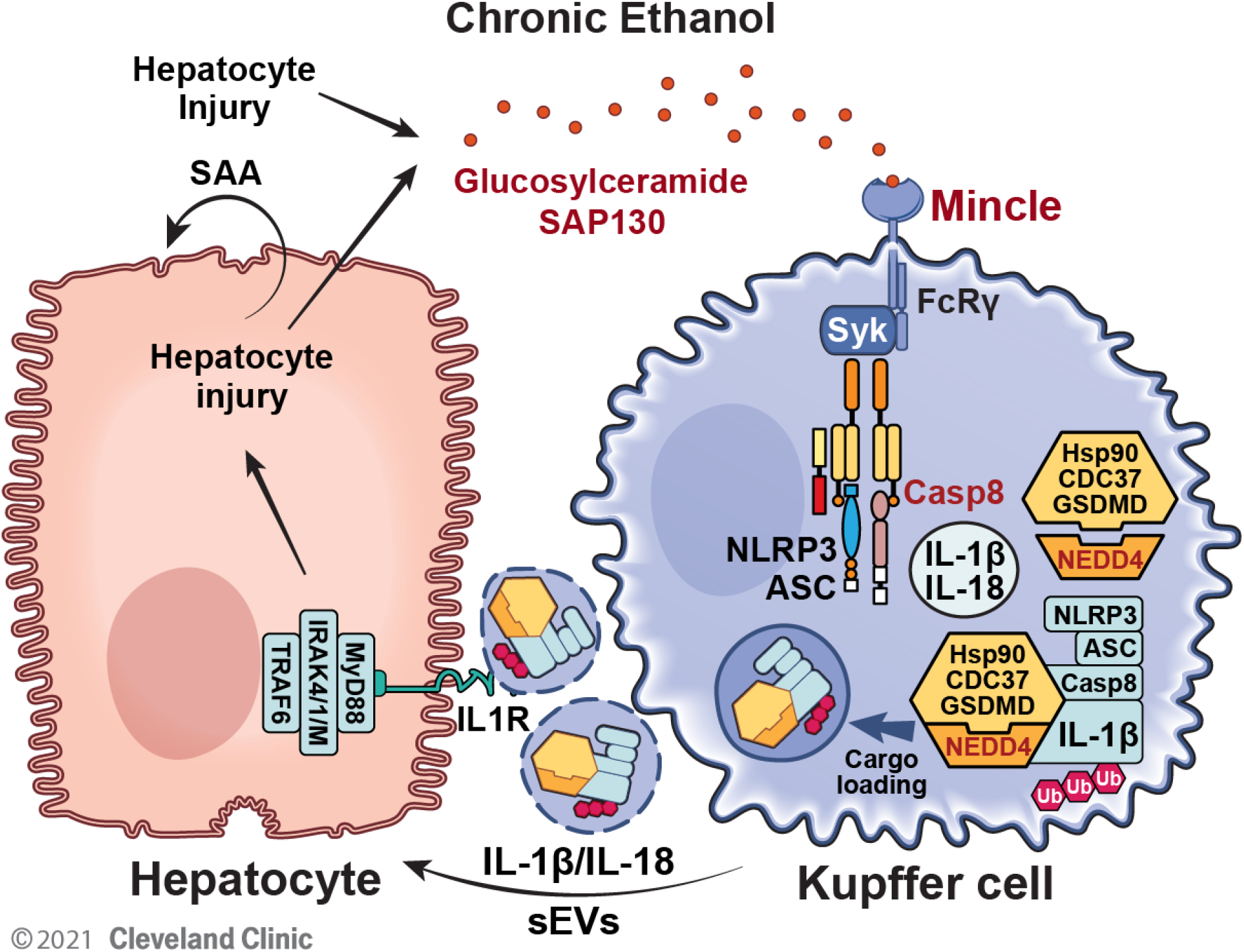
Model for the critical link between β-GluCer and the production of IL-1β-containing sEVs. Ethanol exposure results in the release of β-GlucCer from injury hepatocyte. Mincle, expressed by hepatic macrophages, senses β-GluCer and stimulates the release of IL-1β-containing sEVs in a GSDMD-dependent mechanism. The sEVs, in turn, interact with hepatocytes to stimulate the expression of the acute phase protein SAA. The Mincle-GSDMD-IL-1β pathway provides a mechanism linking hepatocyte injury to inflammation to perpetuate chronic inflammation in ALD.

The concentration of β-GluCer is tightly regulated and restricted to endoplasmic reticulum and Golgi apparatus in normal living cells. Elevated circulating β-GluCer concentrations are observed in various human diseases, including Gaucher disease, multiple sclerosis, and non-alcoholic fatty liver disease (NAFLD). Gaucher disease is an inherited genetic disorder caused by mutation of a critical GluCer hydrolysis enzyme (glucocerebrosidase), leading to accumulation of GluCer in multiple organs, including the liver (40). The fact that liver disease is common in Gaucher disease (41), implicates the critical pathogenic impact of GluCer on hepatic cellular function. In support of this, a recent study indicated that GluCer accumulation in sphingomyelin synthase 1-deficient mouse liver resulted in steatosis, steatohepatitis and fibrosis (42). Consistently, pharmaceutic inhibition of glucosylceramide synthase alleviated the hepatic steatosis and fibrosis in obese mice (43). Collectively, the current study, in combination with previous work, identify β-GluCer, a danger signal released by damaged cells, as a potent pathogenic mediator in driving the progression of liver disease. Out study, in particular, identifies a critical link between β-GluCer and the production of pathogenic IL-1β-containing sEVs that could be an important target for the development of future therapeutic strategies for the prevention or treatment of ALD, as well as liver diseases of other etiologies.

## Abbreviations

Mincle: macrophage inducible c-type lectin
GSDMD: gasdermin d
IL-1β: interleukin-1β
β-GluCer: β-glucosylceramide
LPS: lipopolysaccharide
ALD: alcohol liver disease
sEV: small extracellular vesicles
AH: alcohol-associated hepatitis
Casp: caspase
CDC37: cell deivision cycle 37
HSP90: heat shock protein 90
NEDD4: neural precursor cell expressed developmentally down-regulated protein 4
SAP130: spliceosome-associated protein 130
NLRP3: NLR family pyrin domain containing 3
MELD: model for end-stage liver disease
TDB: trehalose-6,6-dibehenate
ATP: adenosine triphosphate
DAMP: Damage-associated molecular pattern
HC: Healthy control
LDH: Lactate Dehydrogenase
imKC: immortalized Kupffer cell
SAA1: Serum Amyloid A1
ALT: alanine aminotransferase
AST: aspartate aminotransferase
TNFa: tumour necrosis factor alpha
WT: wild type
EtOH: ethanol.

## Acknowledgements

We sincerely thank Dr. Zhaoli Sun at Johns Hopkins University for the human liver specimens.

## Author contributions

Q.Z performed experiments, and wrote the manuscript. W.L., K.B., H.W., M.R.M., R.Z. and X.W. performed experiments. N.W., J.D., and S.D. collected patient samples. L.E.N. and X.L. initiated the idea and wrote the manuscript.

