## Supplemental files for "Mincle-GSDMD-mediated release of IL-1β containing small extracellular vesicles contributes to ethanol-induced liver injury"

**Supplemental Experimental Procedures**

***Mouse model***

All mice were on C57BL/6 background. *Mincle* deficient, *Gsdmd* deficient and *Gsdmd^fl/fl^* (Cyagen Biosciences) mice were previously described (1-3). Wild type mice were purchased from Jackson Laboratories. All procedures involving animals were approved by the Cleveland Clinic Institutional Animal Care and Use Committee. Ten- to twelve-week-old female knock out and littermate mice were housed two per micro-isolator cage and were maintained in a temperature regulated facility with a 12 hour light-dark cycle, with Nylabones provided for environmental enrichment. For Gao-binge model, mice were allowed free access to a Lieber-DeCarli liquid diet (Dyets, Bethlehem, PA; Cat#710260) containing ethanol at 5% (v/v) (28% of total calories) or pair-fed a control diet that isocalorically substituted maltose dextrin for ethanol for 10 days. On the final day of the experiment, pair-fed mice were gavaged with 5g/kg maltose and ethanol-fed mice were gavaged with 5 g/kg ethanol in water. Mice were euthanized 6 h after gavage. Blood was transferred to EDTA-containing tubes for the isolation of plasma. Plasma was then stored at -80°C. The livers were perfused, excised and sectioned for analysis of RNA, protein and histology. Portions of the liver were minced for explant culture.

***Cell and liver explant culture***

Primary hepatocytes and primary Kupffer cells from mice were isolated and cultured as previous described (3, 4). The immortalized mouse Kupffer cell line (imKC) was purchased from Sigma (Cat#SCC119). Primary Kupffer cells and imKCs were primed with 100 pg/ml LPS (Sigma, L2630) overnight followed by stimulation with 20ug/ml GluCer (Avanti, 860539), 2ug/ml TDB (Invivogen, tlrl-tdb) for 12 hrs, or 2.5mM ATP (Sigma, A2383) for 1hr. Whole cell lysates and culture medium were collected for western blot analysis. Cell culture media were used for ELISA, isolation of exosomes (EVs) or cell cytotoxicity analysis. Primary hepatocytes were cultured with ethanol (150 mM), recombinant IL-1β (10 ng/ml) or sEVs isolated from imKCs treated with GluCer + EtOH for 36h. Cytotoxicity was measured by LDH assay kit according to manufacturer’s instructions. For liver explant culture, mouse livers were minced (~2mm) and cultured overnight in serum-free culture medium (DMEM). Culture media were collected and used for isolation of EVs.

***Generation of knockdown and restored cell lines.***

To generate *Mincle* and *Gsdmd* knockdown (KD) cells, control and targeting shRNAs (Mincle, (Origene, TL510331) or Gsdmd (Origene, TG510919)) were delivered to imKC cells by lentiviral infection. Infected cells were cultured in selective medium for 48 hours to enrich for transduced cells before subjected to treatment and analysis. For restoration experiments, HA-tagged WT and D276A mutant of GSDMD were transduced into imKC cells using lentiviral infection.

***Western blotting***

Cells were harvested and lysed in a Triton-containing lysis buffer (0.5% Triton X-100, 20 mM 4-[2-hydroxyethyl]-1-piperazine ethanesulfonic acid [pH 7.4], 150 mM NaCl, 12.5 mM β-glycerophosphate, 1.5 mM MgCl_2_, 10 mM sodium fluoride, 2 mM dithiothreitol, 1 mM sodium orthovanadate, 2 mM ethylene glycol tetraacetic acid, 1 mM phenylmethylsulfonyl fluoride, and complete protease inhibitor cocktail; Roche). For westerns on cell culture medium, protein from culture medium was precipitated by mixing cell-free supernatants with methanol and chloroform at a 4:4:1 ratio. Precipitates were collected by centrifugation and washed with methanol. Precipitates were dissolved in 4X Laemmli buffer for Western blot analysis. For western using sEVs, sEVs isolated from an equal volume of cell culture media were loaded in gels; sEVs were not normalized for particle number**.** Proteins were then separated by 10% sodium dodecyl sulfate polyacrylamide gel electrophoresis, transferred to Immobilon-P membranes (Millipore), and subjected to immunoblotting with antibodies against NEDD4 (catalog 2740), Casp8 (catalog 4790), Cdc37 (catalog 4793), NLRP3 (catalog 15101), β-actin (catalog 3700) (all from Cell Signaling Technology), GSDMD (ab209845 and ab219800), CD63 (ab217345), ubiquitin (ab7780) (all from Abcam), Hsp90 (catalog sc-13119; Santa Cruz Biotechnology, Inc.), caspase-1 (catalog AG-20B-0042-C100; AdipoGen Life Sciences), IL-1β (catalog AF-401, R&D Systems; catalog 503505, BioLegend), and IL-18 (catalog 210-401-323; Rockland Immunochemicals, Inc.). Densitometric analyses of western blots were performed using ImageJ software. Representative western blots with densitometric analysis of three independently performed experiments are shown.

***Co-immunoprecipitation.***

IL-1β co-immunoprecipitation was performed as previously described (1). Briefly, culture medium or sEVs were lysed with NP40 containing lysis buffer (50mM Tris-HCl (pH 7.4), 150mM NaCl, 1%NP-40 and 5mM EDTA and incubated overnight at 4°C with biotinylated hamster anti-mouse IL-1β antibody (BioLegend, 503501). The next day, neutravidin agarose (Pierce Biotechnology) was added to each sample. After 1 hour of incubation at 4°C, agarose resins were centrifuged, washed 3 times in lysis buffer and heated in 30–40 μL 2X Laemmli buffer for SDS-PAGE.

***Quantitative real-time PCR.***

Total RNA was isolated using TRIzol reagent (Invitrogen). 2μg of total RNA was then used for reverse transcription reaction using SuperScript-reverse transcriptase (Invitrogen). RT-PCR was performed in AB 7300 RealTime PCR System, and the gene expression of murine IL-6, TNFα, IL-1β, SAA1, 18s RNA and GAPDH was measured by SYBR® GREEN PCR Master Mix (Applied Biosystems). The results were normalized with the housekeeping gene β-actin or GAPDH.

***ELISA assay.***

IL-1β concentration in cell-free culture medium collected from cell cultures or exosomes from liver explant culture was measured with Duoset ELISA kits (R&D system) following the manufacturer’s instructions.

***Glucosylceramides (GluCer) extraction and quantification***

*GluCer species* from patient serum and mouse plasma were extracted with lipid extraction kit (Biovision). The extracted *GluCer* were analyzed using the HPLC online tandem mass spectrometry method (LC-MS/MS) (23, 45). The compound β-GluCer (d18:1/12:0) was used as internal standard for the calibration of all the GluCer species in the samples.

***ALT/AST and triglyceride measurement***

Plasma samples were assayed for alanine aminotransferase (ALT) and aspartate aminotransferase (AST) (Sekisui Diagnostics, Lexington, MA) following the manufacturer’s instructions. Total hepatic triglycerides were assayed with the Triglyceride Reagent Kit (Cayman Chemical, 10010303) following the manufacturer’s instructions.

***Transmission electron microscopy.***

For ultrastructural analysis, isolated exosomes were fixed for 1 hour at room temperature in 2% paraformaldehyde, 2.5% glutaraldehyde (Polysciences), and 0.05% malachite green (MilliporeSigma) in 100 mM sodium cacodylate buffer, pH 7.2. Malachite green was incorporated into the fixative for stabilization of lipid constituents soluble in aqueous glutaraldehyde. Samples were washed in cacodylate buffer and postfixed for 1 hour in 1% osmium tetroxide (Polysciences). Samples were then rinsed extensively in distilled water before en bloc staining for 1 hour with 1% aqueous uranyl acetate (Ted Pella). Following several rinses in distilled water, samples were dehydrated in a graded series of ethanol and embedded in Eponate 12 resin (Ted Pella). Sections 95 nm in thickness were cut with a Leica Ultracut UC7 ultramicrotome (Leica Microsystems), stained with uranyl acetate and lead citrate, and viewed on a Tecnai G2 Spirit BioTWIN Transmission Electron Microscope (FEI Co.) at 60 kV.

**Supplementary Figure 1.**

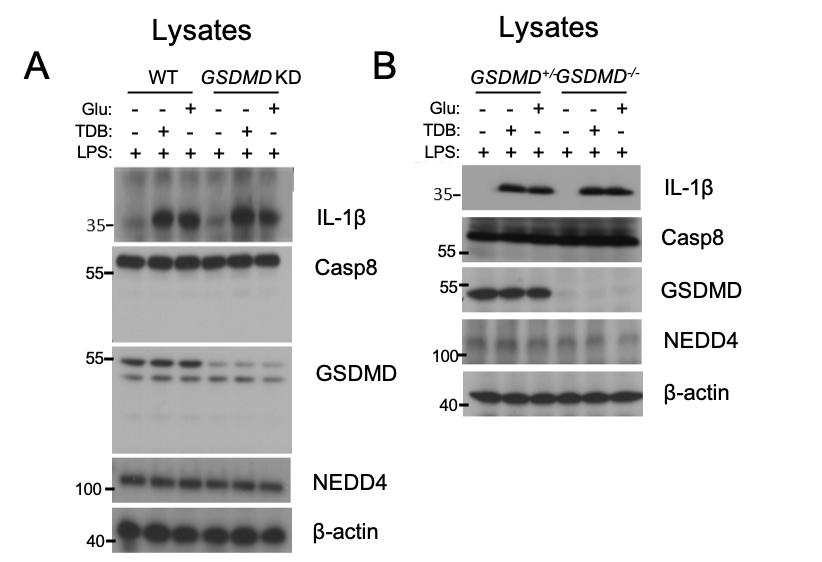

**Supplementary Figure 1: *Gsdmd* deficiency did not impact expression of intracellular IL-1β or components of the sEVs**

(**A,B**) Western blot analysis of indicated protein in the cell lysates from *Gsdmd*-KD imKC **(Fig. 3C**) and Kupffer cells from *Gsdmd*^-/-^ mice (**Fig. 3E)**.

**Supplementary Figure 2.**
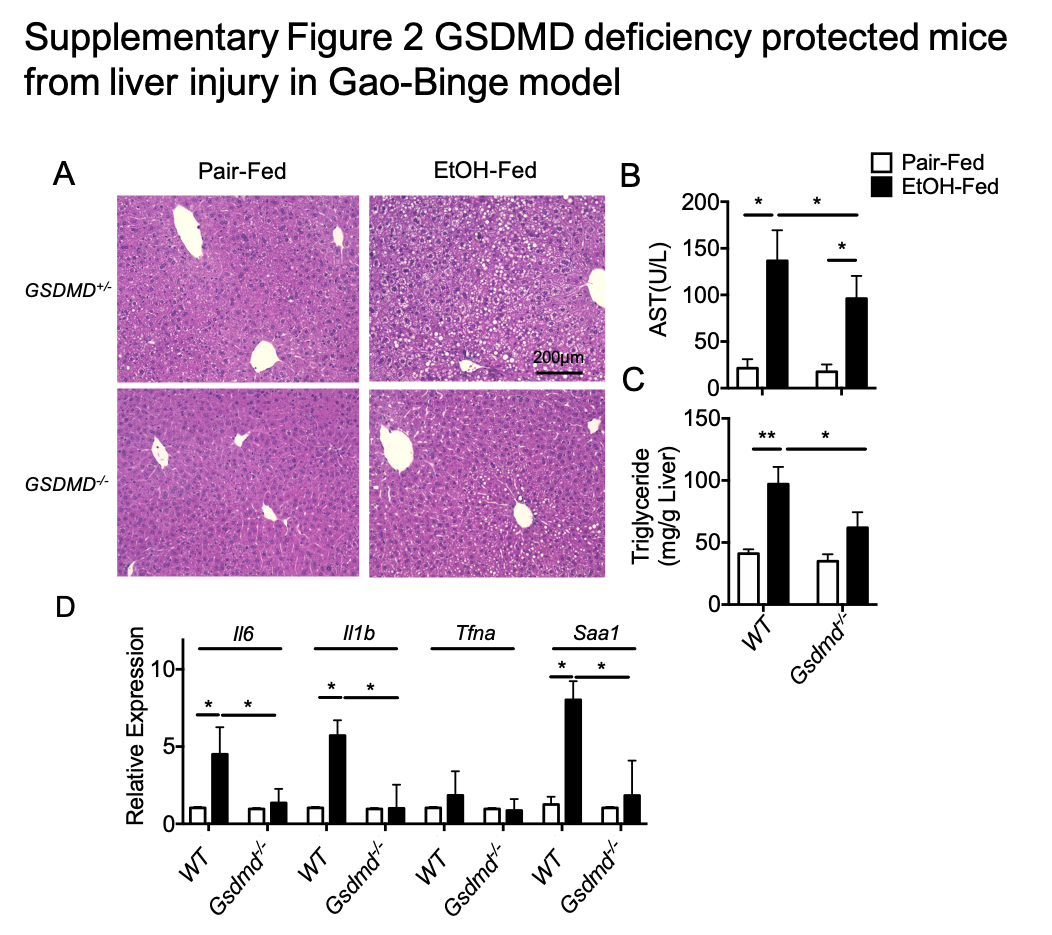

**Supplementary Figure 2:** ***Gsdmd-*deficiency protected mice from liver injury in Gao-Binge model of ethanol-induced liver injury**

*Gsdmd^+/-^* and *Gsdmd^-/-^* mice (n=6) were exposed to Gao-binge ethanol feeding. (**A**) H&E staining for liver histology. (**B**) AST activity in plasma. **(C)** Hepatic triglyceride content. (**D**) IL-6, TNFα, IL-1β and SAA1 mRNA expression. Data represent mean ± SEM. ANOVA. *P<0.05, **P<0.01,

**Supplementary Table 1. Demographic and clinical data for healthy controls and patients with AH (samples used in Fig. 1).**

|  | ASH Mod | ASH Severe | Alcoholic Cirrhosis | Healthy control |
| --- | --- | --- | --- | --- |
| Total Number of Subjects Enrolled | 6 | 9 | 9 | 8 |
| Age (Mean and SD, Range) | 49 ± 3 | 43 ± 9 | 50 ± 9 | 48 ± 12 |
| Gender (N, %) |  |  |  |  |
| Male | 3 (50) | 2 (22) | 6 (67) | 4 (50) |
| Female | 3 (50) | 7 (78) | 3 (33) | 4 (50) |
| Race (N, %) |  |  |  |  |
| White | 5 (83) | 8 (89) | 8 (89) | 7 (88) |
| African American | 1 (16) | 0 | 1 (11) | 0 |
| Asian/Asian American | 0 | 1 (11) | 0 | 1 (12) |
| Other | 0 | 0 | 0 | 0 |
| Ethnicity (N, %) |  |  |  |  |
| Hispanic or Latino | 0 | 1 (11) | 0 | 0 |
| Not Hispanic or Latino | 6 (100) | 8 (89) | 8 (100) | 8 (100) |
| Unknown | 0 | 0 | 0 | 0 |
| Work Status (N, %) |  |  |  |  |
| Unemployed | 5 (83) | 2 (22) | 4 (45) | 5 (63) |
| Employed | 1 (16) | 3 (33) | 2 (22) | 3 (38) |
| Unknown | 0 | 4 (45) | 3 (33) | 0 |

|  | ASH Mod | ASH Severe | Alcoholic Cirrhosis | Healthy control |
| --- | --- | --- | --- | --- |
| Albumin (g/dL) | 3.7 ± 0.6 (2.7-4.3) | 2.6 ± 0.4 (1.7-3.0) | 3.6 ±1.1 (1.9-5.1) | 4.2 ± 0.2 (4.0-4.7) |
| Alkaline Phosphatase (U/L) | 132 ± 47 (76-222) | 211 ± 100 (46-336) | 120 ± 37 (50-171) | 73 ±23 (34-111) |
| ALT (U/L) | 53 ± 70 (9-209) | 39 ± 18 (20-74) | 27 ± 13 (15-54) | 22 ± 8 (12-36) |
| AST (U/L) | 40 ± 14 (27-65) | 118 ± 49 (44-200) | 48 ± 19 (23-83) | 22 ± 6 (16-34) |
| Bilirubin (mg/dL) | 1.7 ± 2.1 (0.2-1.7) | 13.2 ± 4.6 (8.5-20) | 7.3 ± 12.0 (0.3-32.3) | 0.9 ± 0.7 (0.2-2.4) |
| BMI (kg/m2) | 25 ± 4 (19-30) | 27 ± 4 ( 22-36) | 24 ± 7 (16-39) | 27 ± 3 (25-30) |
| Creatinine (mg/dL) | 0.8 ± 0.2 (0.6-1.1) | 1.1 ± 1.2 (0.4-4.4) | 1.2 ± 0.9 (0.7-3.6) | 0.9 ± 0.2 (0.7-1.2) |
| MELD | 9 ± 4 (6-18) | 26 ± 5 (21-34) | 13 ± 14 (7-21) | Not done |
| INR | 1.1 ± 0.2   (1.0-1.5) | 1.7 ± 0.2   (1.3-2.1) | 1.5 ± 0.4  (1.1-2.2) | Not done |
| Total Protein (g/dL) | 7.0 ± 1.2 (6.1-8.3) | 6.2 ± 0.8 (4.9-7.2) | 7.4 ± 1.3 (6.2-9.3) | 7.1 ± 0.4 (6.7-7.2) |

Values reported as mean ± standard deviation (range)

**Supplementary Table 2. Primers used in this study**

| **Primers used in RT-qPCR** | |
| --- | --- |
| **Name** | **sequence (5’ > 3’)** |
| mGapdh | F: CATCACTGCCACCCAGAAGACTG |
|  | R: ATGCCAGTGAGCTTCCCGTTCAG |
| mIL-6 | F: GAGGATACCACTCCCAACAGACC |
|  | R: AAGTGCATCATCGTTGTTCATACA |
| mIL-1β | F: GCAACTGTTCCTGAACTCAACT |
|  | R: ATCTTTTGGGGTCCGTCAACT |
| mTNFα | F: CGTCAGCCGATTTGCTATCT |
|  | R: CGGACTCCGCAAAGTCTAAG |
| mSAA1 | F: ACACCAGGATGAAGCTACTCACCA |
|  | R: CCCTTGGAAAGCCTCGTGAACAAA |
